# Stuck on you! Social brain stimulation increases the cognitive effort required to return to the egocentric perspective

**DOI:** 10.1101/2025.08.26.671988

**Authors:** D. Eliášová, M. Gorelik, A. Chevalier, R. Sherlock, W. Che, M. Meinzer, A.K. Martin

**Affiliations:** School of Psychology, The University of Kent, Canterbury, UK; Department of Neurology, University Medicine Greifswald, Greifswald, Germany; Kent Medway Medical School, Canterbury, UK

**Keywords:** switching, social cognition, social brain, visual perspective taking, non-invasive brain stimulation

## Abstract

Flexible switching between self and other perspectives is critical for adaptive social cognition and is thought to rely on the dynamic regulation of self–other representations. Although neuroimaging implicates the dorsomedial prefrontal cortex (dmPFC) and right temporoparietal junction (rTPJ) in perspective-taking, causal evidence for their specific contributions to perspective switching is lacking. Here, we applied focal transcranial direct current stimulation (f-tDCS) to the dmPFC and rTPJ while participants completed a visual perspective-taking task requiring switches between egocentric and altercentric viewpoints. Anodal stimulation to either site selectively increased the cognitive cost of switching back to the egocentric-perspective, without affecting switches into the altercentric-perspective. Rather than facilitating re-engagement with self-referential processing, stimulation enhanced altercentric persistence or impaired disengagement from the altercentric perspective. These findings provide novel causal evidence that both the dmPFC and rTPJ are involved in regulating the inhibition and updating of self–other representations during perspective switching. Results suggest that stimulation of these hubs may disrupt efficient realignment to the self, highlighting their role in maintaining an altercentric cognitive state. Future studies are required to uncover the precise neural computations that account for the comparable behavioural outcomes observed across distinct social brain hubs.

---

The ability to flexibly switch between one’s own perspective and that of another is a cornerstone of social cognition, underpinning everyday communication, cooperation, and conflict resolution. Yet despite its importance, the cognitive and neural mechanisms that enable perspective switching remain incompletely understood. Behavioural research shows that representing both viewpoints and shifting between them places heavy demands on executive control, but it is not clear how different brain regions contribute to these processes, or how they interact to resolve conflicts between self and other. Neuroimaging work has repeatedly highlighted the right temporoparietal junction (rTPJ) and dorsomedial prefrontal cortex (dmPFC) as key hubs (Healey & Grossman, 2018; Schurz et al., 2015), but causal evidence is sparse. Brain stimulation studies point to a functional dissociation—rTPJ supporting the adoption of another’s perspective, and dmPFC regulating egocentric bias (Martin, Huang, et al., 2019)—yet the causal role of these regions in perspective switching is untested. Clarifying these mechanisms is not only a theoretical challenge for cognitive neuroscience, but also carries translational significance: practitioners working with conditions marked by social difficulties (e.g., autism, psychosis, brain injury) could benefit from mechanistic insights into how perspective-taking succeeds or fails, and how it might be strengthened through targeted interventions.

Visual perspective taking refers to the ability to understand what another person can see or how they see it (Kessler & Rutherford, 2010). It is typically measured using spatial tasks that require participants to judge a scene from another person’s viewpoint. One common paradigm is the Director Task (Apperly et al., 2010), in which participants follow instructions from a “director” who has a limited view of the objects, requiring them to inhibit their own perspective. This task exemplifies level-one perspective taking, which involves understanding or representing what another person can or cannot see. Perspective-taking ability can be assessed by comparing performance when taking another’s perspective versus one’s own and has been used to demonstrate an egocentric bias in both communication (Keysar et al., 2000) and social cognition (Martin, Perceval, et al., 2019). Alternatively, the influence of a task-irrelevant perspective can be investigated by comparing performance between congruent and incongruent trials, with incongruent trials slowing responses due to interference (Martin, Perceval, et al., 2019). However, the cost of switching between perspectives provides a further metric, offering insight into the cognitive demands of maintaining and flexibly shifting between perspectives.

Switching between perspectives places substantial demands on cognitive resources. When self and other perspectives diverge, participants exhibit larger congruency effects— slower responses when the two perspectives conflict—and show increased fixations on task-relevant targets, as revealed by eye-tracking (Ferguson et al., 2017). Elevated fixation counts are widely interpreted as an index of heightened cognitive demand, reflecting the need for additional attentional resources to extract and integrate information. This demonstrates a switch-dependent cognitive demand related to perspective selection. Moreover, asymmetrical switching costs have been demonstrated. For example, Samuel et al. (2019) found a cost of switching when returning to take their own perspective, but not when switching to adopt another person’s point of view. This was taken as evidence that greater inhibition is required to inhibit the egocentric perspective and lifting this inhibition incurs a greater processing cost. Their findings suggest that the egocentric-perspective operates as the cognitive default, necessitating active suppression when taking another’s perspective. Once inhibited, reactivating the egocentric-perspective requires additional executive control, reflected in increased response times and error rates. These results underscore the role of executive processes, particularly inhibition and cognitive control, in managing perspective shifts.

Despite perspective switching being a fundamental process underpinning higher-order social cognition, including theory of mind (ToM; Bradford et al., 2019) and referential communication (Damen et al., 2019), little is known about underlying brain-behaviour associations. The dmPFC is thought to play a key role in managing and integrating social information, including maintaining distinct self–other representations and flexibly selecting between them (Denny et al., 2012; Martin et al., 2017; Wittmann et al., 2016, 2021). In functional imaging studies, activity in this region also increases when tasks require greater cognitive control or a switching of strategy (Clairis & Lopez-Persem, 2023). On the contrary, the rTPJ has been closely linked to the ability to inhibit one’s own egocentric perspective and to simulate another’s viewpoint (Martin, Huang, et al., 2019; Payne & Tsakiris, 2017; Soutschek et al., 2016). It is especially active during tasks that require adopting a non-egocentric perspective, including higher-order social cognitive tasks (Schurz et al., 2014a). Together, these regions form part of a broader social-cognitive network that enables the dynamic and flexible adoption of different visual perspectives in complex social contexts.

Non-invasive brain stimulation (NIBS) offers a method for moving from correlational to causal evidence in the study of social cognition. One such technique, focal transcranial direct current stimulation (f-tDCS), involves modulating activity in specific brain regions through the application of weak electrical currents to the scalp (Meinzer et al., 2024). Previous research has demonstrated the efficacy of tDCS in influencing social cognitive processes (Sellaro et al., 2016), including visual perspective taking (Martin et al., 2017, 2020, 2021; Martin, Huang, et al., 2019; Martin, Su, et al., 2019; Van Elk et al., 2017; Yao et al., 2021). Across multiple replicated studies, f-tDCS has provided causal evidence for the involvement of the right temporoparietal junction (rTPJ) and the dorsomedial prefrontal cortex (dmPFC) in modulating the integration and distinction of self–other representations (Martin et al., 2018; Martin, Su, et al., 2019). For instance, Martin et al. (2018) demonstrated dissociable effects of stimulating these two regions on perspective-taking performance. Specifically, stimulation of the dmPFC enhanced the influence of the altercentric perspective during egocentric judgments across both level-one and level-two perspective-taking tasks. In contrast, stimulation of the rTPJ selectively attenuated the influence of the egocentric perspective during altercentric judgments, particularly in tasks requiring embodied perspective-taking. These findings highlight the distinct yet complementary contributions of the dmPFC and rTPJ in regulating self–other representations. However, direct causal evidence for their involvement in dynamic perspective switching remains unknown.

Therefore, the present study used focal transcranial direct current stimulation (f-tDCS) to identify the causal involvement of two key hubs of the social brain, the dorsomedial prefrontal cortex (dmPFC) and the right temporoparietal junction (rTPJ), in perspective switching. Based on prior evidence for their distinct roles in self–other processing, we hypothesize that dmPFC stimulation will increase the cognitive cost of returning to the egocentric-perspective after adopting the altercentric-perspective. This effect is expected to reflect enhanced integration or maintenance of other-related information, making disengagement from the other’s viewpoint more effortful. Conversely, we predict that rTPJ stimulation will reduce the cognitive effort required to switch to the altercentric-perspective, reflecting improved inhibition of the egocentric perspective and more efficient adoption of the altercentric viewpoint. Together, these predictions reflect dissociable contributions of the dmPFC and rTPJ to the dynamic regulation of self–other representations during perspective switching.

## Methods

### Participants

Fifty-two healthy young adults were recruited from [redacted for anonymous review]. All participants were tDCS naive, free from neurological or psychiatric conditions, and reported no substance abuse. All participants provided written consent in accordance with the Declaration of Helsinki (1991: p1194) and completed a safety screening form and brief interview to ensure they were safe to participate. Participants also completed the Hospital Anxiety and Depression Scale (HADS; Zigmond & Snaith, 1983) and the Autism Quotient (AQ; Baron-Cohen et al., 2001). These measures were used to compare groups receiving rTPJ or dmPFC stimulation. All participants were provided with a small monetary compensation or course credits for their time. The Ethics committee of [redacted for anonymous review] approved the study.

### Study Design, Randomization, and Allocation

The study employed a sham-controlled, double-blind, cross-over design. Healthy young adults completed two experimental sessions in which they received either active or sham stimulation, with stimulation site (dmPFC or rTPJ) serving as the between-subjects factor. All participants first completed baseline screening and questionnaires prior to stimulation.

Randomization of stimulation site (dmPFC vs. rTPJ) was performed using a computer-generated randomization list prepared by an investigator who was not otherwise involved in data collection. Stimulation order (active vs. sham) was counterbalanced within each group, such that half of the participants received active stimulation in the first session and half received sham stimulation first. A minimum interval of three days was required between sessions to avoid carry-over effects.

Double-blinding was maintained using the “study mode” of the DC-Stimulator, which requires entry of a stimulation code. Codes were generated and assigned by a researcher independent of recruitment and testing, thereby ensuring that neither participants nor experimenters were aware of stimulation condition during data collection.

### Transcranial Direct Current Stimulation (tDCS)

Stimulation was administered using a one-channel direct current stimulator (DC-Stimulator Plus®, NeuroConn) with two concentric rubber electrodes (Bortoletto, Rodella, Salvador, Miranda, & Miniussi, 2016; Gbadeyan, Steinhauser, McMahon, & Meinzer, 2016; Perceval et al., 2017). The rubber electrodes allow focal stimulation with the conventional one-channel direct current stimulator. The centre electrode was 2.5cm in diameter and the return electrode was 7.5/9.8 cm for both the dmPFC and rTPJ sites. Current modelling for these montages are presented in detail in previous studies (Martin, Huang, Hunold, & Meinzer, 2017; Martin et al., 2018; de Lillo et al., 2025). Electrodes were attached using an adhesive conductive gel (Weaver Ten20® conductive paste) and held firmly in place using an elastic EEG cap placed over the head in a conventional manner whilst covering the electrodes and preventing any electrode movement. The conductive gel was applied in a consistent manner, covering the electrodes with a thickness no more than 1mm. This ensured the electrodes remained in position and prevented the rubber from contacting the skin directly. The gel was prevented from spreading by ensuring the hair was parted such that the gel would contact skin directly and not require excessive pressure. Both dmPFC and rTPJ sites were identified using the 10-20 International EEG system. The dmPFC was located by finding 15% of the distance from the Fz towards the Fpz. The rTPJ was located by finding CP6. For both the sham and anodal stimulation conditions, the current was ramped up to 1 mA over 5 s and then ramped down over 5 s. In the anodal stimulation condition, the current reached 1 mA over 5 s, was maintained at this intensity for 20 min, and was then ramped down over 5 s. In the sham condition, stimulation was delivered for a total of ∼50 s (5 s ramp up, 40 s at 1 mA, 5 s ramp down), after which the device remained on without current for the remainder of the 20 min session to preserve blinding.

Mood was assessed using the Visual Analogue Mood Scale (VAMS; Folstein & Luria, 1973), administered before and after each stimulation session. The VAMS evaluates current emotional states on 0–10 visual analogue scales across eight dimensions: afraid, confused, sad, angry, energetic, tired, happy, and tense. Higher scores reflect greater intensity of the given mood. To quantify change, post-stimulation ratings were subtracted from pre-stimulation ratings. Individual items were further grouped into composite indices: positive mood (energy, happiness) and negative mood (fear, confusion, sadness, anger, fatigue, tension). Composite change scores were included in the analyses.

Adverse effects were assessed after each session using the self-report questionnaire developed by Brunoni et al. (2011). Participants rated the intensity (recoded to: 0 = absent, 1 = mild, 2 = moderate, 3 = severe) and occurrence of common side effects, including headache, neck pain, scalp pain, tingling, itching, burning sensations, skin redness, sleepiness, concentration difficulties, and sudden mood changes. Total scores are presented for analysis.

Finally, to evaluate blinding efficacy, participants were asked at the end of the second session: *“Do you think the active stimulation was delivered in the first session or the second session?”* Participants were forced to select an option even if they were unsure.

### Visual Perspective Taking Task

A detailed description of the Visual Perspective Taking (VPT) Task used in the present study can be found in Martin, Perceval, et al. (2019). Briefly, participants were presented with a street scene containing one to four tennis balls, which appeared randomly across six possible locations. Rubbish bins were strategically placed to render the balls either visible or occluded from the hypothetical gaze of a humanoid avatar, who was positioned at one of three locations along the street. Participants were instructed to consider any tennis balls positioned behind the avatar as not visible from the avatar’s perspective. In a non-agent control block, a traffic light replaced the avatar, and participants were instructed to judge whether the light would directly hit the tennis balls, serving as a non-social analogue of the visual perspective judgment.

Participants were asked to report either how many tennis balls they themselves could see (egocentric perspective) or how many were visible to the avatar, or would be directly hit by the light (altercentric perspective). On half of the trials, the number of visible tennis balls was identical from both egocentric and altercentric viewpoints (congruent trials), while in the other half, the number differed (incongruent trials). The prompt (you or other) remained on the screen for 750ms followed by a fixation cross for 500ms, then the street scene. The scene remained on the screen until the participant made a response (see Figure 1 for a schematic of the VPT task).

**Figure 1.**
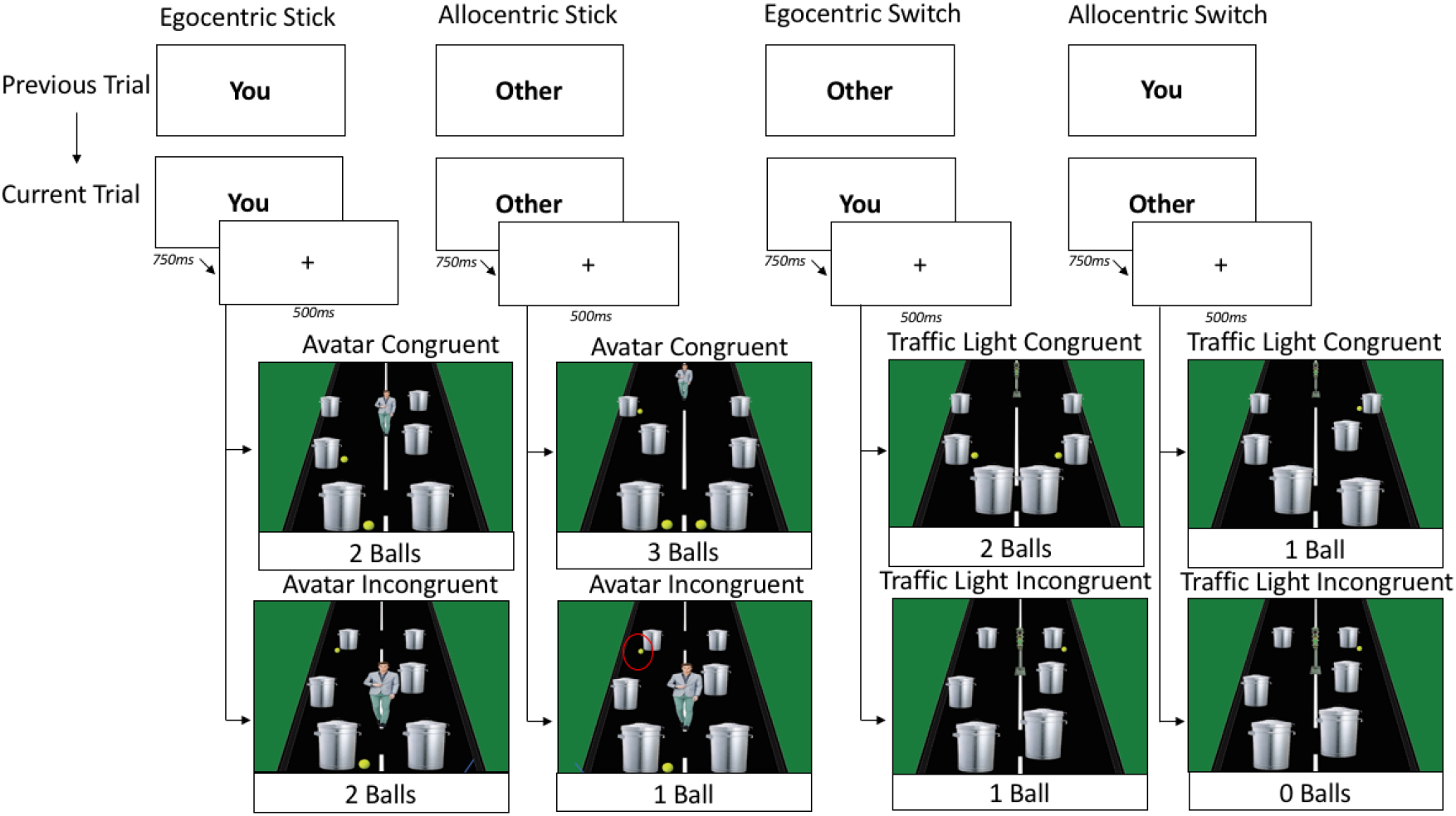
The VPT task (with correct responses for the level one perspective taking task). In the level one perspective taking condition, participants were asked “how many tennis balls can you/other see/light directly shines on?”. If the trial was preceded by the same perspective this was considered a stick trial, whereas if participants were required to change perspectives, this was classified as a switch trial. Only the congruency of the current trial was evaluated in the current study.

On half of the trials, participants were required to maintain the same perspective as on the previous trial (stick trials), while on the other half, they were required to switch perspectives (switch trials). All conditions were fully balanced across the task. The task consisted of four blocks of 88 trials each. The avatar and traffic light blocks were presented separately and completed either in the first two or the final two blocks. Block order was counterbalanced across participants, such that half of the cohort completed the avatar condition first and the other half completed the traffic light condition first. Participants were able to take rest breaks between each block. Participants were informed about the requirements of the new block and presented with ten practice trials for both avatar and light conditions.

The experiment was conducted on desktop computers in dedicated tDCS laboratories at the University of Kent. Stimuli were presented, and responses recorded, using *PsychoPy* (v2023.2). Participants responded via a standard keyboard, with stickers placed over the number keys 1, 2, 9, and 0, corresponding to 1, 2, 3, and 4 tennis balls, respectively. The spacebar was used to indicate 0 tennis balls. Both response latency (in ms) and response accuracy (correct/incorrect) were recorded for each trial. The primary outcome measure was reaction time (RT) for correct responses; accuracy was considered only to exclude incorrect trials from the RT analysis and exclude participants who did not complete the task correctly.

### Statistical Analysis

Response times for correct responses were analysed. Individual responses were removed if they were greater than three standard deviations from the individual’s mean. Participants were excluded from the specific VPT task if they scored 50% correct or less on any condition within the task. This resulted in the removal of 2 participants at the dmPFC stimulation site and 3 at the rTPJ site. All were replaced to achieve the desired sample size. The task was designed to incur minimal response errors. Accuracy was high (>90%) across all conditions and groups and was therefore not analysed in the present study.

All analyses were conducted using JASP version 0.18.3. A 2×2×2×2×2 mixed-design ANOVA was performed with the following within-subjects factors: stimulation (sham vs. anodal), agent (avatar vs. light), congruency (congruent vs. incongruent), and switch (stick vs. switch). Brain region (dmPFC vs. rTPJ) was included as a between-subjects factor. The dependent variable was response time (ms). We present the cognitive effects and stimulation effects separately for ease of interpretation.

Mood change was analysed using two separate 2×2 mixed-design ANOVAS for both negative and positive mood change. The between-subject factor was stimulation site (rTPJ v dmPFC) with stimulation type (sham v anodal) the within-subject factor. Likewise we conducted a 2×2 mixed design ANOVA for total adverse effects with the same factors.

## Results

The stimulation groups (rTPJ v dmPFC) did not differ significantly on age (20.2 vs. 20.9 years), *t*(50) = 0.89, *p* =.38, depressive symptoms (6.0 vs. 7.0), *t*(50) = 0.86, *p* =.40, anxiety symptoms (5.5 vs. 4.8), *t*(50) = 0.59, *p* =.56, or autism quotient (15.9 vs. 17.8), *t*(50) = 1.25, *p* =.22. Group composition was also comparable: each group included 16 women, with 10 men in the rTPJ group and 9 men plus one non-binary participant in the dmPFC group, χ^2^ = 1.05, *p* =.59.

### Cognitive Effects

Independent of stimulation, several robust cognitive effects emerged. A main effect of perspective was observed, *F*(1, 50) = 160.81, *p* <.001, η^2^⍰ =.763, with egocentric-perspective trials (M = 972.68 ms) yielding faster responses than altercentric-perspective trials (M = 1041.49 ms). A strong main effect of congruency was also present, *F*(1, 50) = 264.24, *p* <.001, η^2^⍰ =.841; responses were faster on congruent trials (M = 937.13 ms) than incongruent trials (M = 1077.04 ms). Likewise, there was a significant main effect of switching, *F*(1, 50) = 106.20, *P* <.001, η^2^⍰ =.680, with stick trials (M = 979.01 ms) faster than switch trials (M = 1040.02 ms).

The perspective × congruency interaction was also significant, *F*(1, 50) = 6.64, *P* =.013, η^2^⍰ =.117. The congruency effect was more pronounced under altercentric-perspective trials (congruent: M = 973.12 ms; incongruent: M = 1109.86 ms) than under egocentric-perspective trials (congruent: M = 901.14 ms; incongruent: M = 1042.30 ms; see Figure 2). Finally, a strong congruency × switch interaction was observed, *F*(1, 50) = 119.87, *P* <.001, η^2^⍰ =.706, with the congruency effect amplified in switch trials (congruent: M = 996.46 ms; incongruent: M = 1119.94 ms) compared to non-switch trials (congruent: M = 877.79 ms; incongruent: M = 1034.14 ms; see Figure 3).

**Figure 2.**
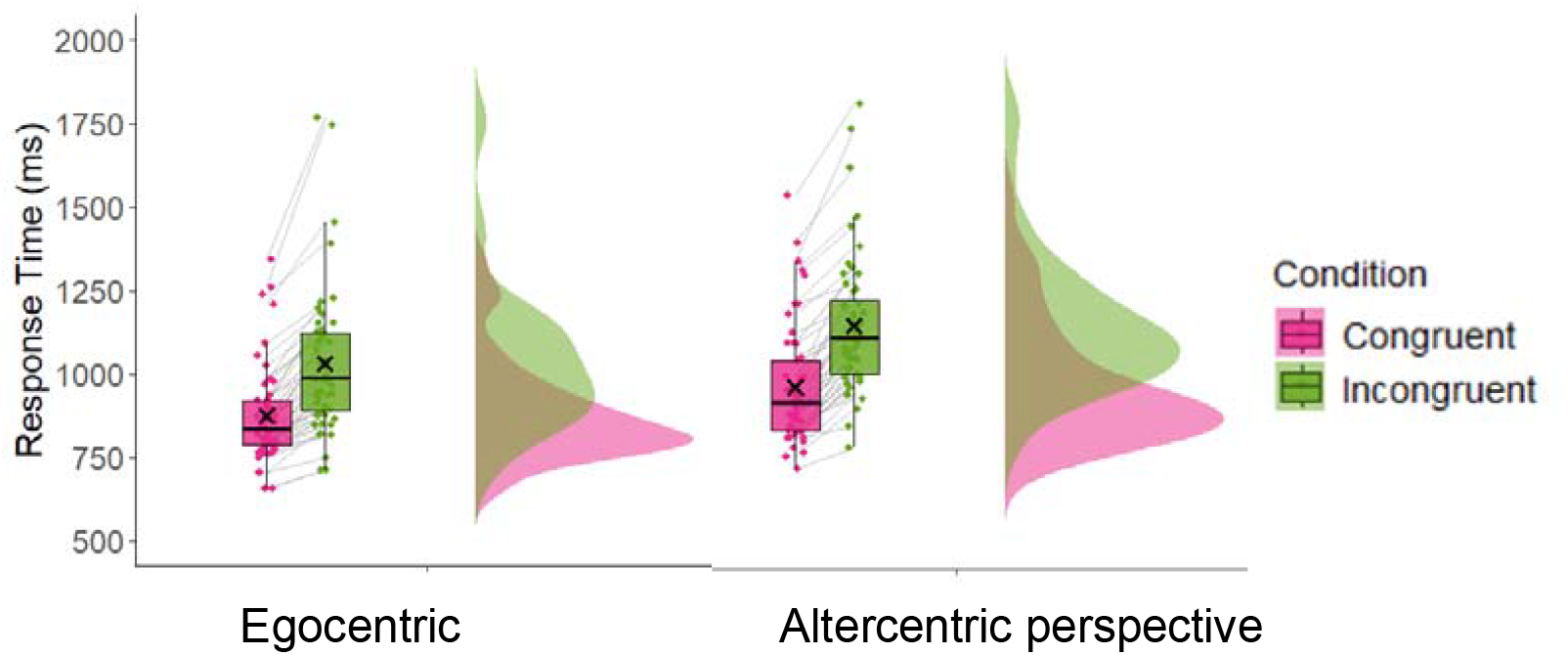
Response times for congruent and incongruent trials during egocentric and altercentric perspective taking.

**Figure 3.**
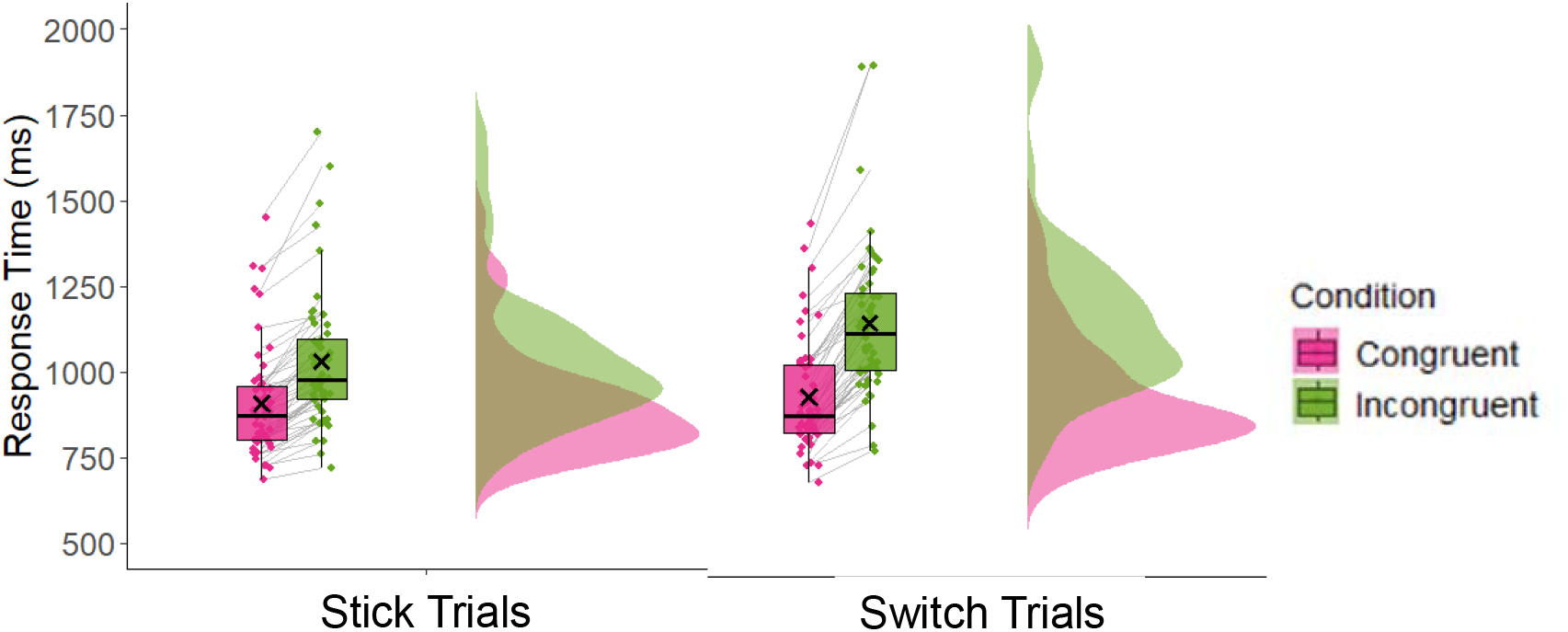
Response times for congruent and incongruent trials during stick or switch trials.

The social nature of the altercentric perspective (avatar v traffic light) did not influence congruency scores. However, the agent × perspective × switch interaction was significant, *F*(1, 50) = 16.94, *P* <.001, η^2^⍰ = 0.25, indicating that the social nature of the altercentric perspective influenced general switching costs. Separate 2×2 RM-ANOVAs were computed for avatar and traffic light conditions. During the avatar condition, the switch x perspective interaction was not significant, F(1,51)=1.21, p=.28, η^2^⍰ = 0.02. However, during the traffic light condition, the switch x perspective interaction was significant, F(1,51)=15.91, p<.001, η^2^⍰ = 0.24. Post-hoc t-tests (Holm corrected), indicated a significant difference between stick and switch trials during egocentric perspective taking, t(51)=6.45, p<.001, d=0.35, but a larger effect during altercentric perspective taking, t(51)=10.87, p<.001, d=0.58 (see Figure 4).

**Figure 4.**
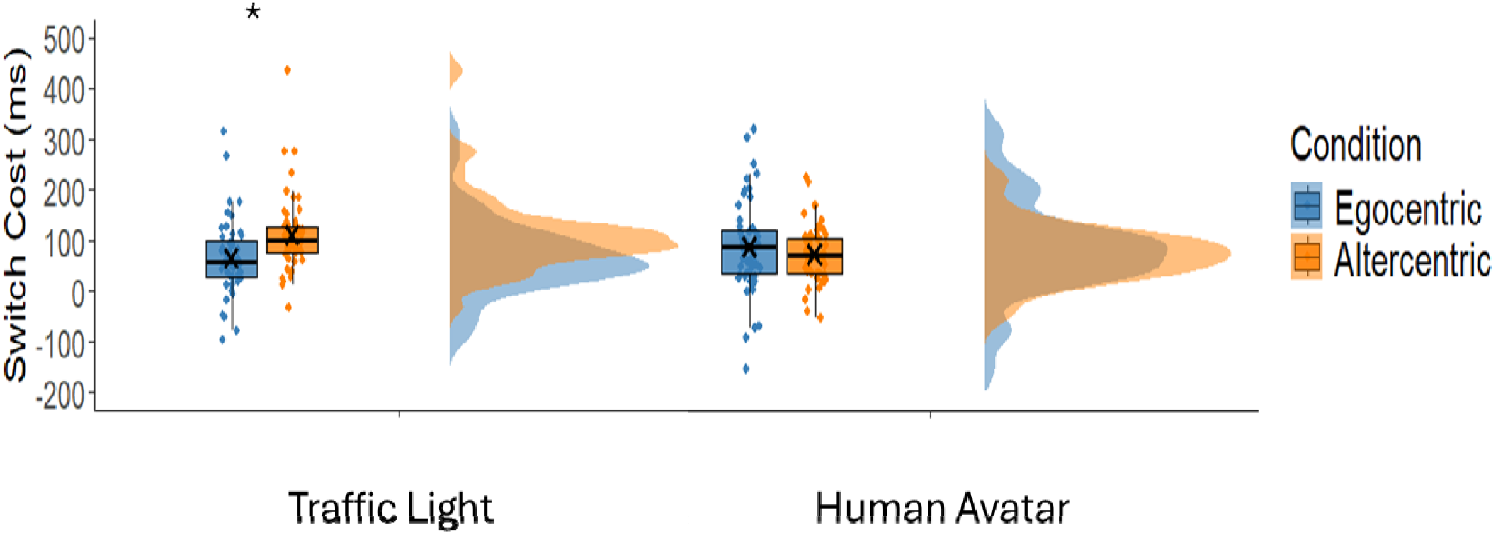
Switch costs when transitioning to or from the egocentric and altercentric perspective for both non-social traffic light and social human avatar conditions

### Stimulation Effects

A significant three-way interaction was identified between stimulation × perspective × switch, *F*(1, 50) = 5.10, *P* =.028, η^2^⍰ = 0.09. However, this did not differ by region, stimulation x perspective x switch x region, *F*(1,50) = 0.45, *P* =.45, η^2^⍰ = 0.01. To explore the stimulation × perspective × switch interaction, separate repeated-measures ANOVAs were conducted for egocentric and altercentric perspective trials.

For egocentric-perspective trials, a significant stimulation × switch interaction was observed, *F*(1, 51) = 4.89, *P* =.032, η^2^⍰ =.09 (see Figure 5.). Post hoc comparisons (Holm-corrected) revealed that during sham stimulation, switch trials (M = 964.41 ms, SE = 30.90) were significantly slower than stick trials (M = 911.21 ms, SE = 28.73), t(51)= 4.89, *P* <.001, *d* = 0.24, but this difference was greater during anodal stimulation, (switch trials: M = 1005.49, SE = 33.29; stick trials: M = 928.63, SE = 28.92), t(51)= 7.09, *P* <.001, *d* = 0.35.

**Figure 5.**
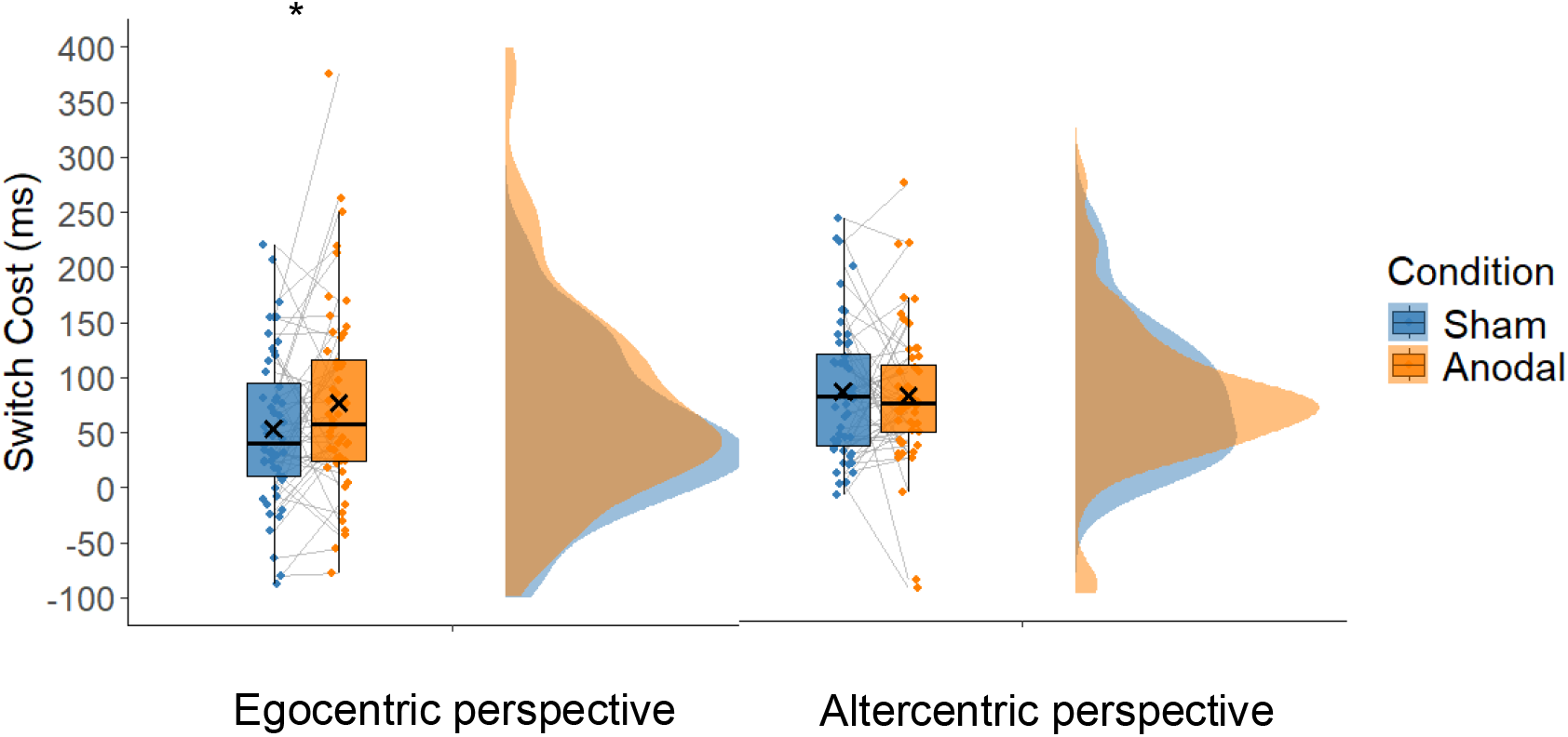
Stimulation to either the dmPFC or the rTPJ increased the cost of switching back to the egocentric perspective.

In contrast, for altercentric-perspective trials, the stimulation × switch interaction was not significant, *F*(1, 51) = 0.15, *P* =.698, η^2^⍰ =.003.

All other higher-order interactions involving stimulation were non-significant (*Ps* >.13). Full statistical output is provided in supplementary table 1.

### Mood change, adverse effects, and blinding

Mood change was minimal after both sham and anodal stimulation sessions (see Table 1). A 2×2 RM-ANOVA examined whether anodal stimulation influenced changes in positive or negative mood from pre-to post-stimulation and whether effects varied by stimulation region. For positive mood, there was no main effect of stimulation, F(1,50)=0.47, p=.50, η^2^⍰=.01, and no interaction with region, F(1,50)=1.65, p=.20, η^2^⍰=.03. Similarly, for negative mood, stimulation had no effect, F(1,50)=0.35, p=.56, η^2^⍰=.01, and no interaction with region, F(1,50)=0.06, p=.81, η^2^⍰<.01.

**Table 1.** Mood change and adverse effects across right TPJ and dmPFC stimulation sites.

|  | rTPJ |  | dmPFC |  |
| --- | --- | --- | --- | --- |
|  | Sham | Anodal | Sham | Anodal |
| Positive Mood | -2.41 (3.42) | -2.07 (3.77) | -0.88 (2.13) | -2.00 (2.60) |
| Negative Mood | 0.02 (4.95) | 0.58 (4.09) | 0.10 (2.00) | 0.34 (3.71) |
| Adverse Effects | 2.65 (2.37) | 3.23 (3.02) | 3.42 (2.23) | 3.46 (2.72) |

Only mild adverse effects were observed (see Table 1). Total adverse effects were not significantly different between sham and active stimulation sessions, *F*(1, 50) = 0.62, *p* =.43, η^2^⍰ = 0.01. No interaction effects were identified for Stimulation × Region, *F*(1, 50) = 0.48, *p* =.49, η^2^⍰ = 0.01.

Blinding was achieved as only 42% (22/52) correctly guessed the active stimulation session overall, 46% at the right TPJ and 38% at the dmPFC which was below chance.

## Discussion

The present study investigated the causal role of the right temporoparietal junction (rTPJ) and dorsomedial prefrontal cortex (dmPFC) in switching between egocentric and altercentric perspectives using focal anodal transcranial direct current stimulation (f-tDCS). Critically, we observed that, contrary to our hypothesis, stimulation to either the rTPJ or dmPFC selectively slowed response times when participants switched from the altercentric to the egocentric perspective, with no such effect observed when switching in the opposite direction. This asymmetric effect provides novel insight into the functional contributions of these regions to the cognitive control of switching during visual perspective taking and the reinstatement of self-referential processing and maintenance of altercentric cognitive processing.

Perspective-taking requires participants to judge visual scenes either from their own point of view (egocentric) or that of another agent (altercentric). Importantly, switching between these perspectives entails the cognitive management of potentially conflicting spatial or visual representations and the inhibition of one perspective in favour of another. The TPJ and dmPFC have been consistently implicated in visual perspective taking, across neuroimaging and brain stimulation studies (Healey & Grossman, 2018; Martin et al., 2017, 2020, 2021; Martin, Huang, et al., 2019; Martin, Su, et al., 2019; Santiesteban et al., 2012; Schurz et al., 2015; Wittmann et al., 2021), yet their distinct, causal contributions to perspective switching had not previously been explored.

Our findings suggest that stimulation to either region interrupts the re-engagement of the egocentric perspective after adopting the viewpoint of another. One possible explanation is that anodal tDCS may have enhanced altercentric processing or increased the salience of the other’s perspective, thereby making it harder to inhibit or disengage when required to return to the egocentric-perspective. This interpretation aligns with prior evidence indicating that the rTPJ and dmPFC support processes such as domain-specific attention reorientation, perspective integration and domain-general conflict monitoring (Corbetta et al., 2008; Gläscher et al., 2012; Martin, Huang, et al., 2019; Oehrn et al., 2014; Yao et al., 2021). Rather than simply adopting another’s viewpoint, these regions appear crucial in resolving self-other representational conflict, a process that becomes especially relevant when transitioning perspectives.

From a mechanistic standpoint, stimulation of the TPJ may have enhanced neural activity related to perspective co-representation, a function widely attributed to this region (Krall et al., 2015; Martin et al., 2021; Martin, Huang, et al., 2019; Santiesteban et al., 2012). By over-activating this system, participants may have experienced greater interference from the altercentric perspective during attempts to re-prioritize their own viewpoint. Similarly, dmPFC stimulation may have amplified processing related to self-other integration or conflict monitoring, potentially increasing deliberation time or cognitive competition between perspectives at the point of switch. However, the lack of a congruency specific effect suggests conflict monitoring or self-other integration is an unlikely explanation. Instead, a more plausible mechanism is that stimulation biased the system toward sustaining the altercentric representation for longer, effectively prioritising the maintenance of another’s perspective. While this may have hindered rapid re-establishment of the egocentric viewpoint in the task, such facilitation of altercentric persistence could reflect a broader role of these regions in supporting flexible social cognition (Schurz et al., 2014b).

Our findings contrast with studies reporting facilitation of perspective-taking following TPJ stimulation (e.g., Santiesteban et al., 2012), though such studies typically focus on the initial adoption of the other’s perspective rather than perspective-switching dynamics. The current results suggest that while TPJ and dmPFC may be essential for adopting another’s viewpoint, their overactivation can hinder the suppression of that viewpoint once it is no longer relevant. This raises important considerations for the use of neuromodulation in enhancing social cognition, as facilitative effects may be context-specific and dependent on the directionality of cognitive control (e.g., switching into vs. out of another’s perspective).

It is also notable that stimulation did not affect switching into the altercentric perspective, suggesting asymmetry in how these regions support perspective transitions. One explanation is that shifting away from the egocentric perspective is typically more cognitively demanding (Martin, Perceval, et al., 2019), requiring suppression of the default self-referential frame in favour of another’s viewpoint. By enhancing activity in the TPJ or dmPFC, regions associated with self-other integration and social perspective-taking, anodal tDCS may promote deeper engagement with the other’s viewpoint. As a result, disengaging from that perspective becomes more effortful, leading to slower reorientation to the self. This interpretation supports the idea that these regions are not simply involved in adopting the other’s perspective, but in sustaining and managing competing representational frames, with stimulation facilitating the persistence of altercentric representations when they are no longer task-relevant.

In addition to the effect of stimulation, several cognitive effects were identified. The observation of greater congruency effects during both switch trials and altercentric trials suggests that cognitive conflict between self and other perspectives is most pronounced when the task demands are highest—namely, when participants must either shift representational frames or adopt a non-default viewpoint, consistent with previous research (Ferguson et al., 2017). Switch trials inherently require reconfiguring the current perspective, which taxes cognitive control mechanisms and may momentarily increase competition between the newly relevant and the recently active perspective. This elevated competition can amplify the interference experienced on incongruent trials, leading to stronger congruency effects. Similarly, altercentric trials demand the suppression of the egocentric-perspective, which is more salient and automatically accessed. As a result, when the other’s viewpoint conflicts with one’s own, the incongruency is felt more acutely, again producing a larger congruency effect. Together, these findings highlight the role of representational conflict and inhibitory control in visual perspective-taking, and suggest that such conflict is modulated not only by the content of the perspectives themselves (i.e., congruent vs. incongruent) but also by the dynamic demands of switching and perspective type.

Importantly, we recognise that the traffic light condition cannot be considered a genuine “perspective” in the same sense as a human avatar. Rather, it serves as a nonsocial analogue in which participants follow a different rule (e.g., “count the dots the light shines on”) instead of representing an alternative viewpoint. From this perspective, the inflated switching costs observed with the traffic light may partly reflect the demands of alternating between two qualitatively different task rules, rather than between two comparable perspectives. Nonetheless, this very asymmetry is informative: it suggests that human avatars provide a more natural anchor for participants, allowing them to treat self and other viewpoints as functionally equivalent and thereby reducing switching demands. In contrast, when the “other” is defined by an arbitrary nonsocial rule, switching becomes more effortful. Thus, while our findings can be interpreted within a rule-switching framework, they also highlight the privileged status of social agents in supporting efficient perspective-shifting. In doing so, our results speak directly to debates about the social nature of VPT tasks (Conway et al., 2017; Santiesteban et al., 2014), showing that the ease of switching is not just about rule complexity but also about whether the task is grounded in a social versus nonsocial context.

The results should be considered in the context of several limitations. The cohort was predominantly female and all were currently engaged in tertiary education in a Western cultural context, with both likely to influence cognition and stimulation effects (Martin, Su, et al., 2019; Wu & Keysar, 2007). Future research should expand to a more generalisable cohort or assess individual differences, both in perspective-switching and the underlying causal neural substrates. The current study focused on switching during level one perspective taking and future research is required to understand whether common or unique effects are associated with switching during more embodied, or level two, perspective taking (Martin et al., 2020). Despite the non-dissociable effects of stimulation to either the dmPFC or the rTPJ, it remains uncertain whether this convergence reflects a shared network-level mechanism or instead the engagement of distinct neural computations that ultimately manifest in comparable behavioural outcomes. On one hand, stimulation to either site may activate a common frontoparietal network supporting self–other distinction, such that effects are propagated across regions irrespective of the precise stimulation locus. On the other hand, the dmPFC may support integration of self-other processes, while the rTPJ may contribute to lower-level attentional reorienting or perspective-taking, yet both can produce overlapping behavioural signatures when excited. Future research will be essential to answer this open question. Combining brain stimulation with neuroimaging approaches such as fMRI or EEG could help to characterise the broader network dynamics evoked by stimulation at each site, revealing whether similar behavioural outcomes arise from shared changes in connectivity or from region-specific patterns of activity. Connectivity analyses, network modelling, and causal manipulations such as dual-site stimulation could further clarify whether the dmPFC and rTPJ make complementary contributions within a distributed system or instead engage separate mechanisms that converge on similar behavioural effects.

In summary, this study provides new insights into the causal role of the rTPJ and dmPFC in perspective switching, highlighting how focal anodal tDCS can interfere with the re-engagement of the egocentric perspective when switching from an altercentric viewpoint. The asymmetric effects on egocentric but not altercentric reorienting suggest that the rTPJ and dmPFC are particularly important in resolving self-other conflicts during egocentric reorienting, where overactivation may disrupt the ability to disengage from the altercentric viewpoint. More broadly, the findings highlight a causal role for these regions in maintaining another’s perspective, a capacity central to social cognition as it underpins processes such as empathy, mentalising, and effective interpersonal communication.

## Acknowledgements

We would like to thank all participants for their time and efforts.

## Conflict of Interest

The authors declare no conflict of interests.

## Funding

No funding to report.

## Data Availability

Data is available on request.

## Notes

### Competing Interest Statement

The authors have declared no competing interest.

## References

Apperly, I. A., Carroll, D. J., Samson, D., Humphreys, G. W., Qureshi, A., & Moffitt, G. (2010). Why are there limits on theory of mind use? Evidence from adults’ ability to follow instructions from an ignorant speaker. Quarterly Journal of Experimental Psychology, 63(6), 1201–1217. 10.1080/17470210903281582

Baron-Cohen, S., Wheelwright, S., Skinner, R., Martin, J., & Clubley, E. (2001). The autism-spectrum quotient (AQ): Evidence from Asperger syndrome/high-functioning autism, males and females, scientists and mathematicians. Journal of Autism and Developmental Disorders, 31(1), 5–17. 10.1023/a:1005653411471

Bradford, E. E. F., Gomez, J.-C., & Jentzsch, I. (2019). Exploring the role of self/other perspective-shifting in theory of mind with behavioural and EEG measures. Social Neuroscience, 14(5), 530–544. 10.1080/17470919.2018.1514324

Brunoni, A. R., Amadera, J., Berbel, B., Volz, M. S., Rizzerio, B. G., & Fregni, F. (2011). A systematic review on reporting and assessment of adverse effects associated with transcranial direct current stimulation. International Journal of Neuropsychopharmacology, 14(8), 1133–1145. 10.1017/S1461145710001690

Clairis, N., & Lopez-Persem, A. (2023). Debates on the dorsomedial prefrontal/dorsal anterior cingulate cortex: Insights for future research. Brain, 146(12), 4826–4844. 10.1093/brain/awad263

Conway, J. R., Lee, D., Ojaghi, M., Catmur, C., & Bird, G. (2017). Submentalizing or mentalizing in a Level 1 perspective-taking task: A cloak and goggles test. Journal of Experimental Psychology: Human Perception and Performance, 43(3), 454–465. 10.1037/xhp0000319

Corbetta, M., Patel, G., & Shulman, G. L. (2008). The Reorienting System of the Human Brain: From Environment to Theory of Mind. Neuron, 58(3), 306–324. 10.1016/j.neuron.2008.04.017

Damen, D., van Amelsvoort, M., van der Wijst, P., & Krahmer, E. (2019). Changing views: The effect of explicit perception-focus instructions on perspective-taking. Journal of Cognitive Psychology, 31(3), 353–369. 10.1080/20445911.2019.1606000

de Lillo, M. D., Korpal, A., Ferguson, H., & Martin, A. K. (2025). A Causal and Dissociable Role for the Right Inferior Prefrontal Cortex in Empathy for Physical and Social Pain (p. 2025.01.19.633761). bioRxiv. 10.1101/2025.01.19.633761

Denny, B. T., Kober, H., Wager, T. D., & Ochsner, K. N. (2012). A meta-analysis of functional neuroimaging studies of self- and other judgments reveals a spatial gradient for mentalizing in medial prefrontal cortex. Journal of Cognitive Neuroscience, 24(8), 1742–1752. 10.1162/jocn_a_00233

Ferguson, H. J., Apperly, I., & Cane, J. E. (2017). Eye tracking reveals the cost of switching between self and other perspectives in a visual perspective-taking task. Quarterly Journal of Experimental Psychology (2006), 70(8), 1646–1660. 10.1080/17470218.2016.1199716

Gläscher, J., Adolphs, R., Damasio, H., Bechara, A., Rudrauf, D., Calamia, M., Paul, L. K., & Tranel, D. (2012). Lesion mapping of cognitive control and value-based decision making in the prefrontal cortex. Proceedings of the National Academy of Sciences, 109(36), 14681–14686. 10.1073/pnas.1206608109

Healey, M. L., & Grossman, M. (2018). Cognitive and Affective Perspective-Taking: Evidence for Shared and Dissociable Anatomical Substrates. Frontiers in Neurology, 9, 491. 10.3389/fneur.2018.00491

Kessler, K., & Rutherford, H. (2010). The Two Forms of Visuo-Spatial Perspective Taking are Differently Embodied and Subserve Different Spatial Prepositions. Frontiers in Psychology, 1. 10.3389/fpsyg.2010.00213

Keysar, B., Barr, D. J., Balin, J. A., & Brauner, J. S. (2000). Taking Perspective in Conversation: The Role of Mutual Knowledge in Comprehension. Psychological Science, 11(1), 32– 38. 10.1111/1467-9280.00211

Krall, S. C., Rottschy, C., Oberwelland, E., Bzdok, D., Fox, P. T., Eickhoff, S. B., Fink, G. R., & Konrad, K. (2015). The role of the right temporoparietal junction in attention and social interaction as revealed by ALE meta-analysis. Brain Structure & Function, 220(2), 587–604. 10.1007/s00429-014-0803-z

Martin, A. K., Dzafic, I., Ramdave, S., & Meinzer, M. (2017). Causal evidence for task-specific involvement of the dorsomedial prefrontal cortex in human social cognition. Social Cognitive and Affective Neuroscience, 12(8), 1209–1218. 10.1093/scan/nsx063

Martin, A. K., Huang, J., Hunold, A., & Meinzer, M. (2019). Dissociable Roles Within the Social Brain for Self–Other Processing: A HD-tDCS Study. Cerebral Cortex, 29(8), 3642–3654. 10.1093/cercor/bhy238

Martin, A. K., Kessler, K., Cooke, S., Huang, J., & Meinzer, M. (2020). The Right Temporoparietal Junction Is Causally Associated with Embodied Perspective-taking. The Journal of Neuroscience: The Official Journal of the Society for Neuroscience, 40(15), 3089–3095. 10.1523/JNEUROSCI.2637-19.2020

Martin, A. K., Perceval, G., Davies, I., Su, P., Huang, J., & Meinzer, M. (2019). Visual perspective taking in young and older adults. Journal of Experimental Psychology. General, 148(11), 2006–2026. 10.1037/xge0000584

Martin, A. K., Perceval, G., Roheger, M., Davies, I., & Meinzer, M. (2021). Stimulation of the Social Brain Improves Perspective Selection in Older Adults: A HD-tDCS Study. Cognitive, Affective, & Behavioral Neuroscience, 21(6), 1233–1245. 10.3758/s13415-021-00929-2

Martin, A. K., Su, P., & Meinzer, M. (2019). Common and unique effects of HD-tDCS to the social brain across cultural groups. Neuropsychologia, 133, 107170. 10.1016/j.neuropsychologia.2019.107170

Meinzer, M., Shahbabaie, A., Antonenko, D., Blankenburg, F., Fischer, R., Hartwigsen, G., Nitsche, M. A., Li, S.-C., Thielscher, A., Timmann, D., Waltemath, D., Abdelmotaleb, M., Kocataş, H., Caisachana Guevara, L. M., Batsikadze, G., Grundei, M., Cunha, T., Hayek, D., Turker, S., … Flöel, A. (2024). Investigating the neural mechanisms of transcranial direct current stimulation effects on human cognition: Current issues and potential solutions. Frontiers in Neuroscience, 18. 10.3389/fnins.2024.1389651

Oehrn, C. R., Hanslmayr, S., Fell, J., Deuker, L., Kremers, N. A., Do Lam, A. T., Elger, C. E., & Axmacher, N. (2014). Neural communication patterns underlying conflict detection, resolution, and adaptation. The Journal of Neuroscience: The Official Journal of the Society for Neuroscience, 34(31), 10438–10452. 10.1523/JNEUROSCI.3099-13.2014

Payne, S., & Tsakiris, M. (2017). Anodal transcranial direct current stimulation of right temporoparietal area inhibits self-recognition. Cognitive, Affective & Behavioral Neuroscience, 17(1), 1–8. 10.3758/s13415-016-0461-0

Perceval, G., Martin, A. K., Copland, D. A., Laine, M., & Meinzer, M. (2017). High-definition tDCS of the temporo-parietal cortex enhances access to newly learned words. Scientific Reports, 7(1), Article 1. 10.1038/s41598-017-17279-0

Samuel, S., Roehr-Brackin, K., Jelbert, S., & Clayton, N. S. (2019). Flexible egocentricity: Asymmetric switch costs on a perspective-taking task. Journal of Experimental Psychology: Learning, Memory, and Cognition, 45(2), 213–218. 10.1037/xlm0000582

Santiesteban, I., Banissy, M. J., Catmur, C., & Bird, G. (2012). Enhancing Social Ability by Stimulating Right Temporoparietal Junction. Current Biology, 22(23), 2274–2277. 10.1016/j.cub.2012.10.018

Santiesteban, I., Catmur, C., Hopkins, S. C., Bird, G., & Heyes, C. (2014). Avatars and arrows: Implicit mentalizing or domain-general processing? Journal of Experimental Psychology. Human Perception and Performance, 40(3), 929–937. 10.1037/a0035175

Schurz, M., Kronbichler, M., Weissengruber, S., Surtees, A., Samson, D., & Perner, J. (2015). Clarifying the role of theory of mind areas during visual perspective taking: Issues of spontaneity and domain-specificity. NeuroImage, 117, 386–396. 10.1016/j.neuroimage.2015.04.031

Schurz, M., Radua, J., Aichhorn, M., Richlan, F., & Perner, J. (2014a). Fractionating theory of mind: A meta-analysis of functional brain imaging studies. Neuroscience and Biobehavioral Reviews, 42, 9–34. 10.1016/j.neubiorev.2014.01.009

Schurz, M., Radua, J., Aichhorn, M., Richlan, F., & Perner, J. (2014b). Fractionating theory of mind: A meta-analysis of functional brain imaging studies. Neuroscience and Biobehavioral Reviews, 42, 9–34. 10.1016/j.neubiorev.2014.01.009

Sellaro, R., Nitsche, M. A., & Colzato, L. S. (2016). The stimulated social brain: Effects of transcranial direct current stimulation on social cognition. Annals of the New York Academy of Sciences, 1369(1), 218–239. 10.1111/nyas.13098

Soutschek, A., Ruff, C. C., Strombach, T., Kalenscher, T., & Tobler, P. N. (2016). Brain stimulation reveals crucial role of overcoming self-centeredness in self-control. Science Advances, 2(10), e1600992. 10.1126/sciadv.1600992

Van Elk, M., Duizer, M., Sligte, I., & Van Schie, H. (2017). Transcranial direct current stimulation of the right temporoparietal junction impairs third-person perspective taking. Cognitive, Affective, & Behavioral Neuroscience, 17(1), 9–23. 10.3758/s13415-016-0462-z

Wittmann, M. K., Kolling, N., Faber, N. S., Scholl, J., Nelissen, N., & Rushworth, M. F. S. (2016). Self-Other Mergence in the Frontal Cortex during Cooperation and Competition. Neuron, 91(2), 482–493. 10.1016/j.neuron.2016.06.022

Wittmann, M. K., Trudel, N., Trier, H. A., Klein-Flügge, M. C., Sel, A., Verhagen, L., & Rushworth, M. F. S. (2021). Causal manipulation of self-other mergence in the dorsomedial prefrontal cortex. Neuron, 109(14), 2353-2361.e11. 10.1016/j.neuron.2021.05.027

Wu, S., & Keysar, B. (2007). The effect of culture on perspective taking. Psychological Science, 18(7), 600–606. 10.1111/j.1467-9280.2007.01946.x

Yao, Y.-W., Chopurian, V., Zhang, L., Lamm, C., & Heekeren, H. R. (2021). Effects of non-invasive brain stimulation on visual perspective taking: A meta-analytic study. NeuroImage, 242, 118462. 10.1016/j.neuroimage.2021.118462

