## Supplemental Table 1 for "Stuck on you! Social brain stimulation increases the cognitive effort required to return to the egocentric perspective"

**Supplemtary Table 1. Full statistical output**

**Repeated Measures ANOVA**

| **Within Subjects Effects** | | | | | | | | | | | | |
| --- | --- | --- | --- | --- | --- | --- | --- | --- | --- | --- | --- | --- |
| **Cases** | | **Sum of Squares** | | **df** | | **Mean Square** | | **F** | | **p** | | **η²_p_** |
| stimulation |  | 396227.01268 |  | 1 |  | 396227.01268 |  | 0.66569 |  | 0.41843 |  | 0.01314 |
| stimulation ✻ region |  | 142356.18585 |  | 1 |  | 142356.18585 |  | 0.23917 |  | 0.62695 |  | 0.00476 |
| Residuals |  | 2.97605×10^+7^ |  | 50 |  | 595210.27839 |  |  |  |  |  |  |
| agent |  | 61767.75348 |  | 1 |  | 61767.75348 |  | 1.10334 |  | 0.29858 |  | 0.02159 |
| agent ✻ region |  | 16698.73606 |  | 1 |  | 16698.73606 |  | 0.29828 |  | 0.58739 |  | 0.00593 |
| Residuals |  | 2.79913×10^+6^ |  | 50 |  | 55982.53508 |  |  |  |  |  |  |
| perspective |  | 4.67270×10^+6^ |  | 1 |  | 4.67270×10^+6^ |  | 160.81351 |  | < .001 |  | 0.76282 |
| perspective ✻ region |  | 2144.52793 |  | 1 |  | 2144.52793 |  | 0.07381 |  | 0.78699 |  | 0.00147 |
| Residuals |  | 1.45283×10^+6^ |  | 50 |  | 29056.62808 |  |  |  |  |  |  |
| congruency |  | 1.17048×10^+7^ |  | 1 |  | 1.17048×10^+7^ |  | 264.24269 |  | < .001 |  | 0.84089 |
| congruency ✻ region |  | 16924.03591 |  | 1 |  | 16924.03591 |  | 0.38207 |  | 0.53930 |  | 0.00758 |
| Residuals |  | 2.21478×10^+6^ |  | 50 |  | 44295.57147 |  |  |  |  |  |  |
| switch |  | 2.22551×10^+6^ |  | 1 |  | 2.22551×10^+6^ |  | 106.19670 |  | < .001 |  | 0.67989 |
| switch ✻ region |  | 1199.47613 |  | 1 |  | 1199.47613 |  | 0.05724 |  | 0.81190 |  | 0.00114 |
| Residuals |  | 1.04783×10^+6^ |  | 50 |  | 20956.50797 |  |  |  |  |  |  |
| stimulation ✻ agent |  | 4068.70229 |  | 1 |  | 4068.70229 |  | 0.12001 |  | 0.73048 |  | 0.00239 |
| stimulation ✻ agent ✻ region |  | 22881.67721 |  | 1 |  | 22881.67721 |  | 0.67492 |  | 0.41524 |  | 0.01332 |
| Residuals |  | 1.69515×10^+6^ |  | 50 |  | 33903.00446 |  |  |  |  |  |  |
| stimulation ✻ perspective |  | 1078.89835 |  | 1 |  | 1078.89835 |  | 0.04122 |  | 0.83994 |  | 0.00082 |
| stimulation ✻ perspective ✻ region |  | 53505.24636 |  | 1 |  | 53505.24636 |  | 2.04403 |  | 0.15902 |  | 0.03928 |
| Residuals |  | 1.30882×10^+6^ |  | 50 |  | 26176.31787 |  |  |  |  |  |  |
| agent ✻ perspective |  | 5303.20243 |  | 1 |  | 5303.20243 |  | 0.46228 |  | 0.49970 |  | 0.00916 |
| agent ✻ perspective ✻ region |  | 5160.11504 |  | 1 |  | 5160.11504 |  | 0.44981 |  | 0.50551 |  | 0.00892 |
| Residuals |  | 573589.91636 |  | 50 |  | 11471.79833 |  |  |  |  |  |  |
| stimulation ✻ congruency |  | 17561.54455 |  | 1 |  | 17561.54455 |  | 1.04799 |  | 0.31090 |  | 0.02053 |
| stimulation ✻ congruency ✻ region |  | 2.51692 |  | 1 |  | 2.51692 |  | 0.00015 |  | 0.99027 |  | 3.00396×10^-6^ |
| Residuals |  | 837864.95263 |  | 50 |  | 16757.29905 |  |  |  |  |  |  |
| agent ✻ congruency |  | 7060.04788 |  | 1 |  | 7060.04788 |  | 0.80850 |  | 0.37287 |  | 0.01591 |
| agent ✻ congruency ✻ region |  | 948.60445 |  | 1 |  | 948.60445 |  | 0.10863 |  | 0.74308 |  | 0.00217 |
| Residuals |  | 436615.38732 |  | 50 |  | 8732.30775 |  |  |  |  |  |  |
| perspective ✻ congruency |  | 75790.15418 |  | 1 |  | 75790.15418 |  | 6.63650 |  | 0.01299 |  | 0.11718 |
| perspective ✻ congruency ✻ region |  | 195.67892 |  | 1 |  | 195.67892 |  | 0.01713 |  | 0.89638 |  | 0.00034 |
| Residuals |  | 571010.18346 |  | 50 |  | 11420.20367 |  |  |  |  |  |  |
| stimulation ✻ switch |  | 10168.94898 |  | 1 |  | 10168.94898 |  | 1.39704 |  | 0.24281 |  | 0.02718 |
| stimulation ✻ switch ✻ region |  | 6131.69022 |  | 1 |  | 6131.69022 |  | 0.84239 |  | 0.36312 |  | 0.01657 |
| Residuals |  | 363946.57707 |  | 50 |  | 7278.93154 |  |  |  |  |  |  |
| agent ✻ switch |  | 103539.68609 |  | 1 |  | 103539.68609 |  | 14.82403 |  | < .001 |  | 0.22868 |
| agent ✻ switch ✻ region |  | 2753.93742 |  | 1 |  | 2753.93742 |  | 0.39429 |  | 0.53291 |  | 0.00782 |
| Residuals |  | 349229.15772 |  | 50 |  | 6984.58315 |  |  |  |  |  |  |
| perspective ✻ switch |  | 27392.40006 |  | 1 |  | 27392.40006 |  | 3.10544 |  | 0.08414 |  | 0.05848 |
| perspective ✻ switch ✻ region |  | 406.72058 |  | 1 |  | 406.72058 |  | 0.04611 |  | 0.83085 |  | 0.00092 |
| Residuals |  | 441038.83910 |  | 50 |  | 8820.77678 |  |  |  |  |  |  |
| congruency ✻ switch |  | 776391.96941 |  | 1 |  | 776391.96941 |  | 119.87458 |  | < .001 |  | 0.70567 |
| congruency ✻ switch ✻ region |  | 5782.72606 |  | 1 |  | 5782.72606 |  | 0.89285 |  | 0.34925 |  | 0.01754 |
| Residuals |  | 323835.12694 |  | 50 |  | 6476.70254 |  |  |  |  |  |  |
| stimulation ✻ agent ✻ perspective |  | 11815.62307 |  | 1 |  | 11815.62307 |  | 2.36059 |  | 0.13074 |  | 0.04508 |
| stimulation ✻ agent ✻ perspective ✻ region |  | 9328.75659 |  | 1 |  | 9328.75659 |  | 1.86375 |  | 0.17830 |  | 0.03594 |
| Residuals |  | 250268.42930 |  | 50 |  | 5005.36859 |  |  |  |  |  |  |
| stimulation ✻ agent ✻ congruency |  | 4645.63768 |  | 1 |  | 4645.63768 |  | 1.25722 |  | 0.26754 |  | 0.02453 |
| stimulation ✻ agent ✻ congruency ✻ region |  | 6426.00048 |  | 1 |  | 6426.00048 |  | 1.73902 |  | 0.19327 |  | 0.03361 |
| Residuals |  | 184758.92871 |  | 50 |  | 3695.17857 |  |  |  |  |  |  |
| stimulation ✻ perspective ✻ congruency |  | 5413.14305 |  | 1 |  | 5413.14305 |  | 0.52762 |  | 0.47100 |  | 0.01044 |
| stimulation ✻ perspective ✻ congruency ✻ region |  | 62.59796 |  | 1 |  | 62.59796 |  | 0.00610 |  | 0.93805 |  | 0.00012 |
| Residuals |  | 512978.23981 |  | 50 |  | 10259.56480 |  |  |  |  |  |  |
| agent ✻ perspective ✻ congruency |  | 10656.80663 |  | 1 |  | 10656.80663 |  | 1.24219 |  | 0.27038 |  | 0.02424 |
| agent ✻ perspective ✻ congruency ✻ region |  | 3070.92990 |  | 1 |  | 3070.92990 |  | 0.35796 |  | 0.55234 |  | 0.00711 |
| Residuals |  | 428952.65182 |  | 50 |  | 8579.05304 |  |  |  |  |  |  |
| stimulation ✻ agent ✻ switch |  | 127.97809 |  | 1 |  | 127.97809 |  | 0.02113 |  | 0.88500 |  | 0.00042 |
| stimulation ✻ agent ✻ switch ✻ region |  | 11.96596 |  | 1 |  | 11.96596 |  | 0.00198 |  | 0.96472 |  | 0.00004 |
| Residuals |  | 302773.73533 |  | 50 |  | 6055.47471 |  |  |  |  |  |  |
| stimulation ✻ perspective ✻ switch |  | 19709.52212 |  | 1 |  | 19709.52212 |  | 5.10251 |  | 0.02828 |  | 0.09260 |
| stimulation ✻ perspective ✻ switch ✻ region |  | 2706.67437 |  | 1 |  | 2706.67437 |  | 0.70072 |  | 0.40653 |  | 0.01382 |
| Residuals |  | 193135.66400 |  | 50 |  | 3862.71328 |  |  |  |  |  |  |
| agent ✻ perspective ✻ switch |  | 87975.27305 |  | 1 |  | 87975.27305 |  | 16.94067 |  | < .001 |  | 0.25307 |
| agent ✻ perspective ✻ switch ✻ region |  | 3039.36048 |  | 1 |  | 3039.36048 |  | 0.58526 |  | 0.44785 |  | 0.01157 |
| Residuals |  | 259657.05726 |  | 50 |  | 5193.14115 |  |  |  |  |  |  |
| stimulation ✻ congruency ✻ switch |  | 3947.59291 |  | 1 |  | 3947.59291 |  | 0.66008 |  | 0.42038 |  | 0.01303 |
| stimulation ✻ congruency ✻ switch ✻ region |  | 1901.54096 |  | 1 |  | 1901.54096 |  | 0.31796 |  | 0.57536 |  | 0.00632 |
| Residuals |  | 299024.13150 |  | 50 |  | 5980.48263 |  |  |  |  |  |  |
| agent ✻ congruency ✻ switch |  | 28021.75584 |  | 1 |  | 28021.75584 |  | 3.26536 |  | 0.07678 |  | 0.06130 |
| agent ✻ congruency ✻ switch ✻ region |  | 9908.89599 |  | 1 |  | 9908.89599 |  | 1.15468 |  | 0.28773 |  | 0.02257 |
| Residuals |  | 429076.67615 |  | 50 |  | 8581.53352 |  |  |  |  |  |  |
| perspective ✻ congruency ✻ switch |  | 10681.57313 |  | 1 |  | 10681.57313 |  | 2.84933 |  | 0.09764 |  | 0.05391 |
| perspective ✻ congruency ✻ switch ✻ region |  | 1042.96934 |  | 1 |  | 1042.96934 |  | 0.27821 |  | 0.60021 |  | 0.00553 |
| Residuals |  | 187440.29521 |  | 50 |  | 3748.80590 |  |  |  |  |  |  |
| stimulation ✻ agent ✻ perspective ✻ congruency |  | 211.37111 |  | 1 |  | 211.37111 |  | 0.03637 |  | 0.84953 |  | 0.00073 |
| stimulation ✻ agent ✻ perspective ✻ congruency ✻ region |  | 3797.63105 |  | 1 |  | 3797.63105 |  | 0.65339 |  | 0.42273 |  | 0.01290 |
| Residuals |  | 290609.25422 |  | 50 |  | 5812.18508 |  |  |  |  |  |  |
| stimulation ✻ agent ✻ perspective ✻ switch |  | 119.81214 |  | 1 |  | 119.81214 |  | 0.02279 |  | 0.88062 |  | 0.00046 |
| stimulation ✻ agent ✻ perspective ✻ switch ✻ region |  | 1.33455 |  | 1 |  | 1.33455 |  | 0.00025 |  | 0.98735 |  | 5.07650×10^-6^ |
| Residuals |  | 262886.09699 |  | 50 |  | 5257.72194 |  |  |  |  |  |  |
| stimulation ✻ agent ✻ congruency ✻ switch |  | 207.37437 |  | 1 |  | 207.37437 |  | 0.03053 |  | 0.86200 |  | 0.00061 |
| stimulation ✻ agent ✻ congruency ✻ switch ✻ region |  | 3174.14100 |  | 1 |  | 3174.14100 |  | 0.46727 |  | 0.49740 |  | 0.00926 |
| Residuals |  | 339650.09037 |  | 50 |  | 6793.00181 |  |  |  |  |  |  |
| stimulation ✻ perspective ✻ congruency ✻ switch |  | 1669.32249 |  | 1 |  | 1669.32249 |  | 0.28759 |  | 0.59415 |  | 0.00572 |
| stimulation ✻ perspective ✻ congruency ✻ switch ✻ region |  | 827.35989 |  | 1 |  | 827.35989 |  | 0.14254 |  | 0.70737 |  | 0.00284 |
| Residuals |  | 290227.55148 |  | 50 |  | 5804.55103 |  |  |  |  |  |  |
| agent ✻ perspective ✻ congruency ✻ switch |  | 16858.05244 |  | 1 |  | 16858.05244 |  | 3.33195 |  | 0.07392 |  | 0.06248 |
| agent ✻ perspective ✻ congruency ✻ switch ✻ region |  | 6781.15846 |  | 1 |  | 6781.15846 |  | 1.34028 |  | 0.25249 |  | 0.02611 |
| Residuals |  | 252976.13182 |  | 50 |  | 5059.52264 |  |  |  |  |  |  |
| stimulation ✻ agent ✻ perspective ✻ congruency ✻ switch |  | 619.89185 |  | 1 |  | 619.89185 |  | 0.10930 |  | 0.74232 |  | 0.00218 |
| stimulation ✻ agent ✻ perspective ✻ congruency ✻ switch ✻ region |  | 4928.61805 |  | 1 |  | 4928.61805 |  | 0.86903 |  | 0.35570 |  | 0.01708 |
| Residuals |  | 283570.43607 |  | 50 |  | 5671.40872 |  |  |  |  |  |  |
| *Note.*  Type III Sum of Squares | | | | | | | | | | | | |

| **Between Subjects Effects** | | | | | | | | | | | | |
| --- | --- | --- | --- | --- | --- | --- | --- | --- | --- | --- | --- | --- |
| **Cases** | | **Sum of Squares** | | **df** | | **Mean Square** | | **F** | | **p** | | **η²_p_** |
| region |  | 2183.01988 |  | 1 |  | 2183.01988 |  | 0.00205 |  | 0.96405 |  | 0.00004 |
| Residuals |  | 5.32060×10^+7^ |  | 50 |  | 1.06412×10^+6^ |  |  |  |  |  |  |
| *Note.*  Type III Sum of Squares | | | | | | | | | | | | |
